# The mechanosensitive Ca^2+^-permeable ion channel PIEZO1 promotes satellite cell function in skeletal muscle regeneration

**DOI:** 10.1101/2021.03.18.435982

**Authors:** Kotaro Hirano, Masaki Tsuchiya, Seiji Takabayashi, Kohjiro Nagao, Yasuo Kitajima, Yusuke Ono, Keiko Nonomura, Yasuo Mori, Masato Umeda, Yuji Hara

## Abstract

Muscle satellite cells (MuSCs), myogenic stem cells in skeletal muscle, play an essential role in muscle regeneration. During the regeneration process, cues from the surrounding microenvironment are critical for the proliferation and function of MuSCs. However, the mechanism by which mechanical stimuli from the MuSCs niche is converted into biochemical signals to promote muscle regeneration is yet to be determined. Here, we show that PIEZO1, a calcium ion (Ca^2+^)-permeable cation channel that is activated by membrane tension, mediates the spontaneous Ca^2+^ influx to controls the regenerative function of MuSCs. Our genetically engineering approach in mice revealed that PIEZO1 is functionally expressed in MuSCs, and the conditional deletion of *Piezo1* in MuSCs delays myofiber regeneration after myofiber injury, which is at least in part due to the growth defect in MuSCs via the reduction in RhoA-mediated actomyosin formation. Thus, we provide the first evidence in MuSCs that PIEZO1, a *bona fide* mechanosensitive ion channel, promotes the proliferative and regenerative function during skeletal muscle regeneration.

## Introduction

Muscle-resident stem cells called muscle satellite cells (MuSCs) are critical for skeletal muscle regeneration after injuries induced by repeated contraction and relaxation of myofibers. Under a resting condition, quiescent MuSCs (QSCs) reside on the plasma membrane of myofibers and underneath the extracellular matrix [1]. Upon injury, MuSCs are committed to become fusogenic myoblasts to repair the damaged myofibers or to form multinucleated cells called myotubes [2]. The importance of MuSCs is further demonstrated by the fact that its impaired function is closely associated with sarcopenia and a class of muscle diseases (i.e., muscular dystrophy) [3–5].

Extensive efforts have been made to elucidate the mechanisms underlying MuSC-mediated skeletal myofiber regeneration. MuSCs express paired box 7 (Pax7), a transcription factor that is essential for MuSC-specific gene expression [6]. Once activated, the expression of distinct transcription factors, including Myf5 and MyoD, are upregulated, generating myoblast-specific gene signatures [2]. In addition to these stage-specific transcription factors, the fate of MuSCs is also determined by a number of signaling cascades, such as p38/MAPK [7], Notch [8–10], and RhoA/ROCK [11] pathways, during muscle regeneration. Although several secretory molecules (e.g., HGF and Wnt4) have also been identified as regulators of MuSCs [11, 12], the critical determinant that senses the changes in the microenvironmental niche surrounding MuSCs in response to myofiber injuries remains to be elucidated.

The concentration of Ca^2+^ in the cytosol is strictly maintained within the nanomolar range, which is approximately 20,000-fold lower than in extracellular fluid [13]. Thus, Ca^2+^ influx across the plasma membrane is recognized as one of the critical determinants of a variety of cellular and physiological events [14]. Multiple Ca^2+^ channels play critical roles in skeletal muscle functions; L-type voltage-gated Ca^2+^ channel, also known as a dihydropyridine receptor, is essential for myofiber contraction via its interaction with ryanodine receptor [15, 16]. In addition, the STIM1-ORAI complex, a component of store-operated Ca^2+^ channel, is involved in muscle development, growth and physiology [17, 18]. Among Ca^2+^ channels, mechanosensitive ion channels that are activated by physical stimuli toward the plasma membrane have been thought to be plausible candidates as regulators of MuSC functions.

PIEZO1, a mechanosensitive ion channel that is activated by membrane tension, plays a fundamental role in sensing biophysical force [19]. PIEZO1 is well-conserved among species from plants to human, and is composed of approximately 2,500 amino acid residues, forming trimer to act as a mechanosensitive ion channel. Although *PIEZO1* gene mutations have been identified in patients with hereditary xerocytosis [20], studies on tissue-specific *Piezo1*-deficient mice have revealed that PIEZO1 is involved in the mechanosensation of various cells and tissues, including neuronal progenitor cells [21], chondrocytes [22], and blood and lymphatic vessels [23, 24], suggesting that PIEZO1 is critical for tissue homeostasis [25]. We previously reported that PIEZO1 is highly expressed in myoblasts, and that the ion channel activity of PIEZO1 is positively regulated by phospholipid flippase, an enzyme that catalyzes the translocation of phospholipids from the outer to inner leaflets of the plasma membranes. Moreover, our results demonstrated that *Piezo1* deletion led to the formation of abnormally enlarged myotubes due to impaired actomyosin assembly, suggesting that PIEZO1-mediated Ca^2+^ influx is one of critical determinants of morphogenesis during myotube formation [26]. However, the function of PIEZO1 in MuSCs remains to be elucidated.

In this study, we evaluated the expression of PIEZO1 in skeletal muscle using *Piezo1-tdTomato* mice and showed that PIEZO1 is highly expressed in undifferentiated MuSCs. Moreover, *Piezo1* deficiency in MuSCs led to delayed myofiber regeneration after myolysis-induced degeneration, at least in part due to reduced proliferation of MuSCs. Thus, our results revealed that PIEZO1 is involved in skeletal muscle regeneration by promoting the function of MuSCs.

## Results

### PIEZO1 is restrictedly expressed in MuSCs

We previously reported that PIEZO1 is critical for morphogenesis during myotube formation [26]. To examine the expression profile of *Piezo1* in skeletal muscle myogenesis in detail, we isolated the total RNA samples from QSCs, activated muscle satellite cells (ASCs), and mature skeletal muscle tissue. Semi-quantitative RT-PCR analysis demonstrated that *Piezo1* mRNA expression was clearly and moderately detected in QSCs and ASCs, respectively; however, no obvious *Piezo1* expression was observed in mature myofibers, which is consistent with previous literatures, including our previous report (Fig 1A, [19], [26]).

**Figure 1.**
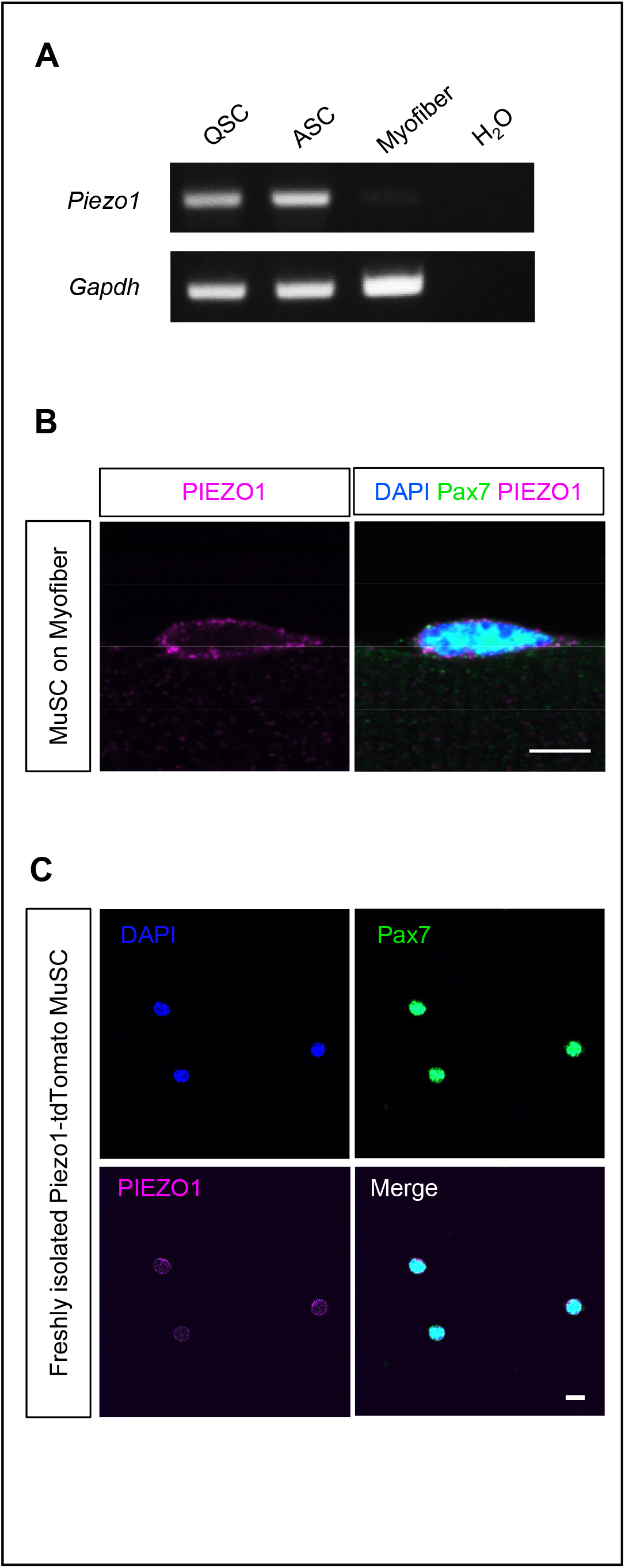
Expression of PIEZO1 channel in undifferentiated MuSCs. **A** Semi-quantitative RT-PCR analysis of the *Piezo1* gene in MuSCs. RNA samples were isolated from QSCs, ASCs, and mature skeletal muscle. *Gapdh* gene was also detected as a control gene. **B** and **C** Detection of PIEZO1 protein in MuSCs. Immunofluorescent analysis of PIEZO1 on isolated myofibers from EDL muscle of *Piezo1-tdTomato* mice (**B**), and on freshly isolated MuSCs from *Piezo1-tdTomato* mice (**C**). PIEZO1-tdTomato was visualized by immunofluorescent staining with anti-RFP antibody (magenta). Pax7 and nuclei were also detected with anti-Pax7 antibody (green) and DAPI (blue), respectively. Bar: 10 μm.

To validate the expression of PIEZO1 in MuSCs at the protein level, we utilized *Piezo1-tdTomato* mice in which the C-terminus of endogenous PIEZO1 was fused with a red fluorescent protein, tdTomato (EV1A, [27]). We performed immunofluorescent analysis to detect PIEZO1 and Pax7, a MuSC-specific transcription factor [6], on floating single myofibers isolated from extensor digitorum longus (EDL) muscle of *Piezo1-tdTomato* mice. PIEZO1 expression was clearly observed in Pax7-positive MuSCs but not in myofibers, which is consistent with the RT-PCR result (Figs 1A and 1B). In addition, PIEZO1 expression was also detected on MuSCs isolated from *Piezo1-tdTomato* muscle samples using fluorescence-activated cell sorting (FACS), as a population of VCAM-1^+ve^, Sca1^−ve^, CD31^−ve^, and CD45^−ve^ cells.

MuSCs are committed to become myoblasts during myogenesis in response to myofiber injury [28]. To investigate whether PIEZO1 is expressed in activated MuSCs, tibialis anterior (TA) muscle was injected with cardiotoxin (CTX), a venom toxin that causes myofiber degeneration and concomitant regeneration, followed by immunofluorescent detection of PIEZO1 on regenerating myofibers after four days. Again, PIEZO1 expression was detected on Pax7-positive cell population (EV1C), suggesting that PIEZO1 could be involved in myogenesis.

### PIEZO1 is involved in Ca^2+^ mobilization in MuSCs

To examine whether PIEZO1 acts as a Ca^2+^-permeable channel in MuSCs, we generated *Piezo1*-deficient mice. As the systemic deletion of the *Piezo1* gene causes embryonic lethality [23], we employed conditional gene targeting using Cre-loxP-mediated genetic recombination. Mice harboring “floxed” alleles of the *Piezo1* gene were crossed with *Pax7^CreERT2/+^*, a transgenic mouse line that specifically expresses Cre-recombinase in MuSCs under the control of tamoxifen (EV2A, [29]). After the resultant mice *Piezo1^flox/flox^*; *Pax7^CreERT2/+^* were obtained, *Piezo1* gene deletion was induced by intraperitoneal injection of tamoxifen to the mice (called *Piezo1* cKO hereafter) for five consecutive days. MuSCs isolated from *Piezo1* cKO were seeded onto glass bottom dishes and were subjected to Ca^2+^ measurements using a ratiometric Ca^2+^ indicator Fura-2. Ca^2+^ measurements demonstrated that Ca^2+^ influx was clearly detected with Yoda-1 (a chemical agonist for PIEZO1 [30]) in MuSCs and myoblasts isolated from wild type but not from *Piezo1* cKO, confirming the PIEZO1 expression and the effectiveness of conditional deletion of *Piezo1* gene in those cells (EV2B and 2C).

Ca^2+^ fluctuations are known to be involved in a variety of cellular processes [14]. We asked if Ca^2+^ fluctuations could be detected in isolated MuSCs. Ca^2+^ measurements clearly detected spontaneous Ca^2+^ transients in HEPES-buffered saline containing 2 mM Ca^2+^ but were almost completely abolished by chelating external Ca^2+^ with EGTA (ethylene glycol-bis(2-aminoethyl ether)-N, N, N’, N’-tetraacetic acid) (Figs 2A and 2B). This result allowed us to examine the role of PIEZO1 in spontaneous Ca^2+^ transients in MuSCs. Indeed, a significant reduction in Ca^2+^ transients was evident in *Piezo1* cKO MuSCs (average value for amplitudes; wild type: 0.0263 ±0.0017, *Piezo1* cKO: 0.0195 ± 0.0015) (Figs 2C-2E). These results indicate that PIEZO1 acts as a Ca^2+^-permeable ion channel that dominantly generates Ca^2+^ fluctuations in MuSCs.

**Figure 2.**
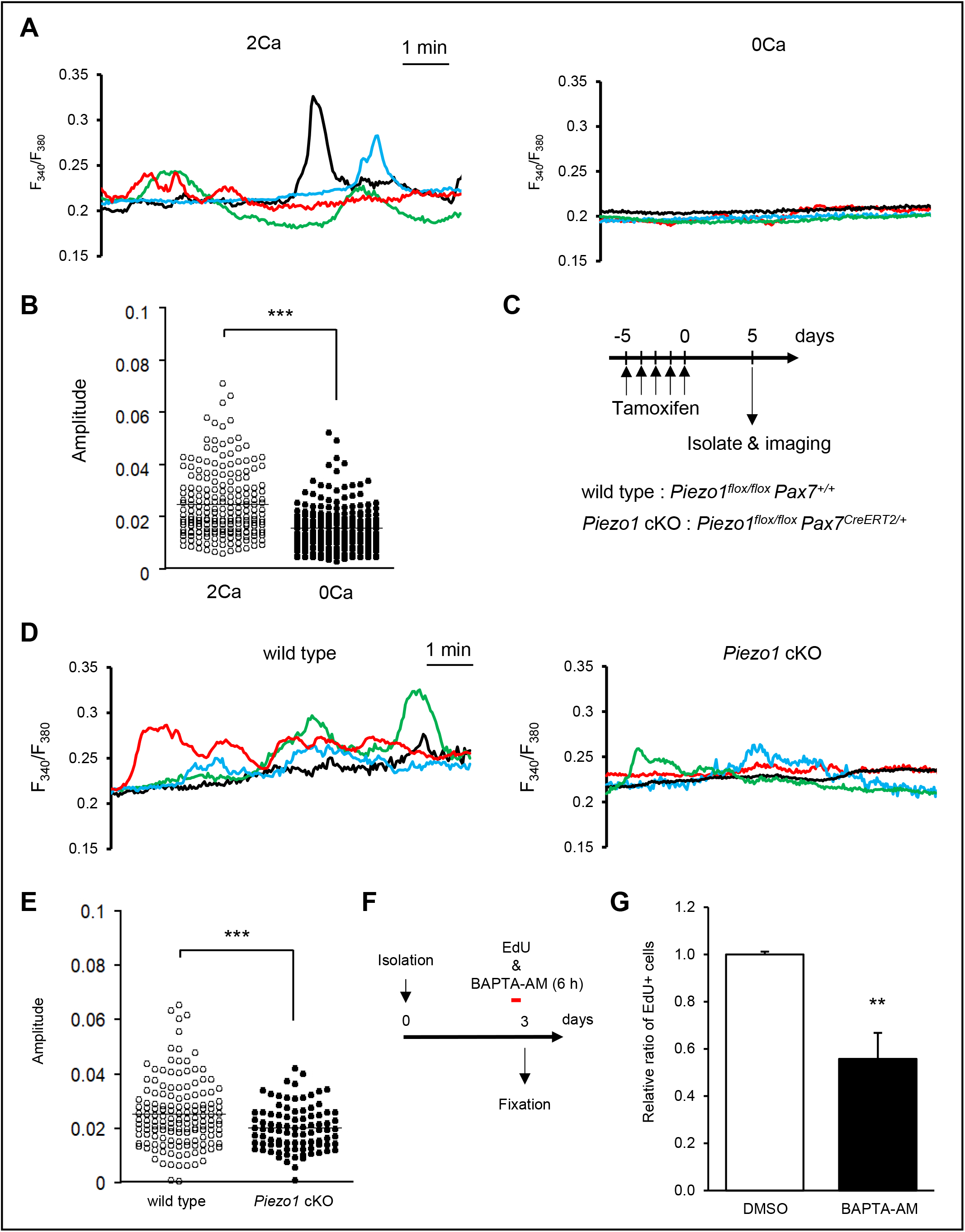
PIEZO1-dependent Ca^2+^ fluctuation in MuSCs. **A** and **B** Calcium ion (Ca^2+^) measurements in MuSCs under a resting condition. After the isolation of MuSCs from wild type mice, Ca^2+^ fluctuations were monitored using a Ca^2+^ indicator fura-2. (**A**) Representative traces of Ca^2+^ fluctuations in wild-type MuSCs in the presence (upper panel) and absence (0 Ca^2+^) of extracellular Ca^2+^. (**B**) Quantification of the amplitude of Ca^2+^ fluctuations in MuSCs. Data represent means ± S.E.M. **C** Time course for induction of *Piezo1*-deficiency. **D and E** Calcium ion (Ca^2+^) measurements in wild type or *Piezo1* cKO MuSCs. (**D**) Representative traces of wild type and *Piezo1* cKO cells were shown in the left and right panels, respectively. (**E**) Quantification of the amplitude of Ca^2+^ fluctuations in MuSCs. Data represent means ± S.E.M. **F** Time course for chelating cytosolic Ca^2+^ in proliferating MuSC. **G** An EdU incorporation assay on cultured MuSCs. The number of EdU+ cells was normalized to that on control; DMSO (N = 4).

Next, we asked whether Ca^2+^ affects the proliferative capacity of MuSCs by chelating cytosolic Ca^2+^ in MuSCs using BAPTA-AM (1,2-bis(2-aminophenoxy) ethane-N,N,N’,N’-tetraacetic acid), a membrane-permeable Ca^2+^ chelator. During the 6-h-pulse of 5-ethynyl-20-deoxyuridine (EdU) with BAPTA-AM (Fig 2F), the number of EdU-positive proliferating MuSCs decreased in BAPTA-AM-treated cells (Fig 2G). This indicates that Ca^2+^ signaling is essential for MuSC proliferation.

### PIEZO1 plays a role in myofiber regeneration

Based on the results of immunofluorescent analyses and Ca^2+^ imaging, we hypothesized that PIEZO1 could play a role in myofiber regeneration. To examine the effect of *Piezo1* deletion on MuSC function, we utilized CTX to induce the degeneration and subsequent regeneration of myofibers (Fig 3A). Reduction in muscle weight was more evident in *Piezo1* cKO mice compared to wild type mice at seven days post-CTX injection (Figs 3A-3C, EV3A and 3B). This result led us to further evaluate the histological abnormalities during myofiber regeneration after CTX injection. Hematoxylin and Eosin staining showed myofibers with centrally located nuclei (i.e., regenerating myofibers) were observed in wild type muscle. In contrast, *Piezo1* cKO muscle displayed a series of abnormalities, such as infiltration of immune cells and fibrosis, which are hallmarks of the defects in myofiber regeneration (Fig 3D). These abnormalities were further confirmed by a reduction in cross-sectional area (CSA) of regenerating myofibers (Fig 3E) and by the increased collagen deposition at seven days post-injection with CTX (Figs 3F and 3G). In addition, heterologous *Piezo1^Tm1a/+^*, a mouse line heterologously harboring the ‘knockout-first’ allele (*Tm1a*) that disrupts the *Piezo1* gene function by inserting gene cassettes expressing *LacZ* and neomycin-resistant genes into the *Piezo1* locus, did not show obvious abnormality (EV4A). Importantly, no obvious histological abnormality was observed in *Piezo1^+/+^*; *Pax7^CreERT2/+^* mice, eliminating the possibility that the heterologous deletion in the *Pax7* gene affects the regeneration capacity of myofibers in *Piezo1* cKO mice, at least in our experimental procedure. These results indicate that PIEZO1 is involved in myofiber regeneration after muscle injury.

**Figure 3.**
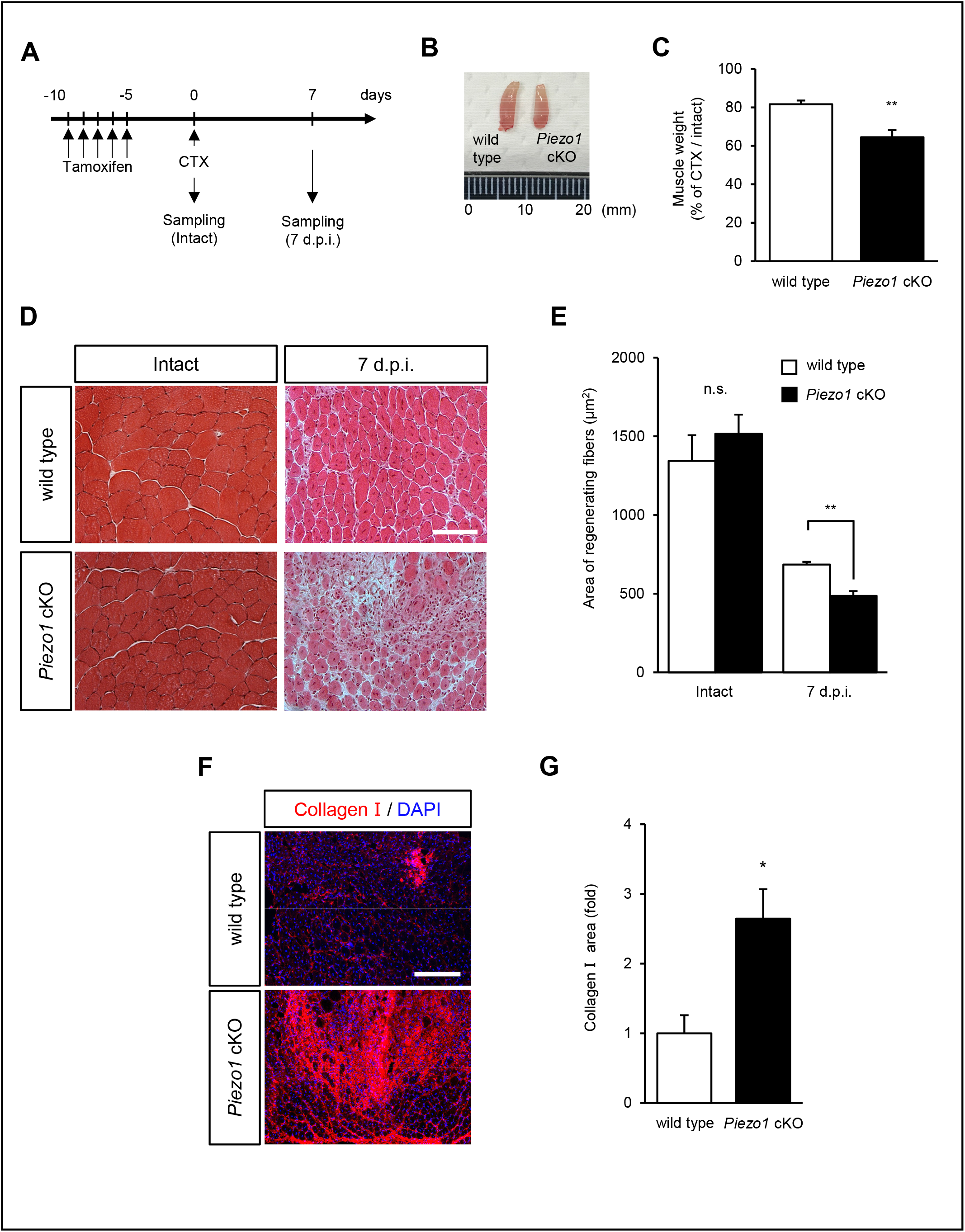
Impaired regeneration capacity of *Piezo1*-deficient muscle after cardiotoxin-induced myofiber degeneration. **A** Time course for the induction of *Piezo1* deficiency, injection of TA with cardiotoxin, and isolation of regenerating muscle samples. **B** Representative images of regenerating TA muscle samples in wild type (left) and *Piezo1* cKO mice. **C** Weight of TA muscle after CTX-induced muscle injury. N = 3–9. **D** Hematoxylin and Eosin staining of cross muscle sections from intact and cardiotoxin-injected TA muscle samples harvested seven days post-injection. Upper panels: wild type, lower panels: *Piezo1* cKO mice. **E** CSA of intact and regenerating myofibers harvested seven days post-cardiotoxin injection. **F** and **G** Fluorescent intensity of regenerating muscle sections stained with anti-collagen I antibody. Red: collagen I, blue: nuclei (DAPI). Bar graphs represent mean ± S.E.M. White bars: wild type, black bars: *Piezo1* cKO muscle. *: *P* < 0.05, **: *P* < 0.01.

### PIEZO1 plays a role in MuSC proliferation

To investigate the role of PIEZO1 during myofiber regeneration, we sought to evaluate the number of MuSCs in CTX-injected TA muscle. Immunofluorescent detection of MuSCs with anti-Pax7 antibody revealed that the number of MuSCs on myofibers isolated from *Piezo1* cKO was significantly reduced compared with that from wild type mice (66.5 ± 3.85 and 46.6 ± 2.47 in wild type and *Piezo1* cKO mice, respectively) (Figs 4A and 4B) seven days after the induction of *Piezo1* deletion. This result suggests that PIEZO1 may play a role in the maintenance of MuSC pool. To further examine whether the proliferation of MuSCs is affected by the *Piezo1* deletion *in vivo*, we performed EdU incorporation assays. After CTX injection to TA muscle, the mice were subjected to intraperitoneal injection with EdU, followed by the detection of incorporated EdU in the nuclei representing active DNA synthesis (Fig 4D). As indicated in Fig 4E, the number of EdU-positive MuSCs was clearly reduced in *Piezo1* cKO muscle sections. Moreover, this trend was also observed in MuSCs cultured for three days (Fig 4F), indicating that PIEZO1 plays a role in activated MuSC proliferation after muscle injury.

**Figure 4.**
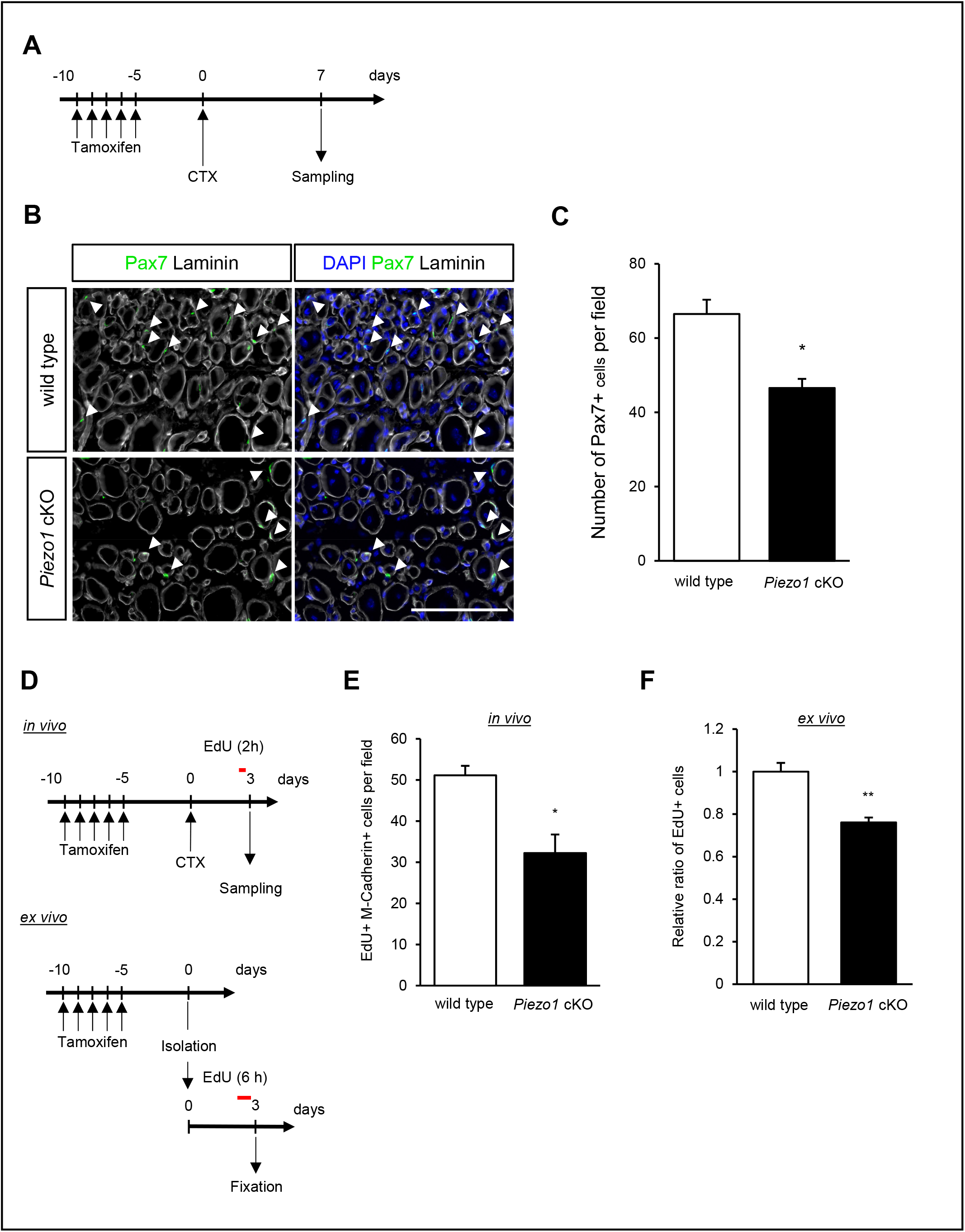
Reduced proliferation capacity of MuSCs by *Piezo1*-deficiency during myofiber regeneration. **A** Time course for induction of *Piezo1* deficiency, cardiotoxin injection, and harvesting myofibers. **B** and **C** Detection of Pax7-positive MuSCs (arrowheads) on cross sections from wild type (left) and *Piezo1* cKO muscle (right). The number of MuSCs was evaluated seven days after TMX treatment. Bar: 100 μm. (**C**) Quantification of the number of Pax7-positive MuSCs per visual field (N = 3). Bar graphs represent mean ± S.E.M. *: *P* < 0.05. **D** Time course for EdU incorporation assays on MuSCs in regenerating TA muscle (*in vivo*) and isolated myofibers (*ex vivo*). **E** EdU incorporation assay on regenerating TA muscle (*in vivo*). After cardiotoxin administration, the number of EdU+ M-Cadherin+ cells (i.e., MuSCs possessing proliferative capacity) was counted in cross sections from wild type and *Piezo1* cKO muscle samples (N = 3). **F** EdU incorporation assay on cultured MuSCs (*ex vivo*). The number of EdU+ cells was normalized to that on wild type (N = 4).

To further evaluate the capacity for myogenic progression, we monitored the expression of MyoD and Myogenin, which are myogenic regulatory factors that are predominantly expressed in MuSCs after activation and proliferation [31], as well as Pax7. The isolated myofibers were cultured in a plating medium for three days, and were subjected to immunofluorescent analysis with co-staining of anti-Pax7 and anti-MyoD or anti-Pax7 and anti-Myogenin antibodies. Our results revealed that, although the number of cells was significantly reduced in *Piezo1* cKO mice, the proportions of Pax7-positive/MyoD-negative (i.e., self-renewed cells), Pax7-positive/MyoD-positive (i.e., activated or proliferative cells), and Pax7-negative/MyoD-positive (i.e., differentiated cells) groups [32] in *Piezo1* cKO mice were comparable to that in wild-type mice (Fig 5C and 5D). The same trend was observed when co-stained with anti-Pax7 and anti-Myogenin, where the proportion of Pax7-positive/Myogenin-negative (i.e., quiescent or activated cells) and Pax7-negative/Myogenin-positive (i.e., differentiated cells) groups in *Piezo1* cKO mice was comparable to that in wild-type mice (Figs 5E and 5F). These results suggest that PIEZO1 plays a role in MuSC proliferation after activation, but not in cell fate decision *in vitro*.

**Figure 5.**
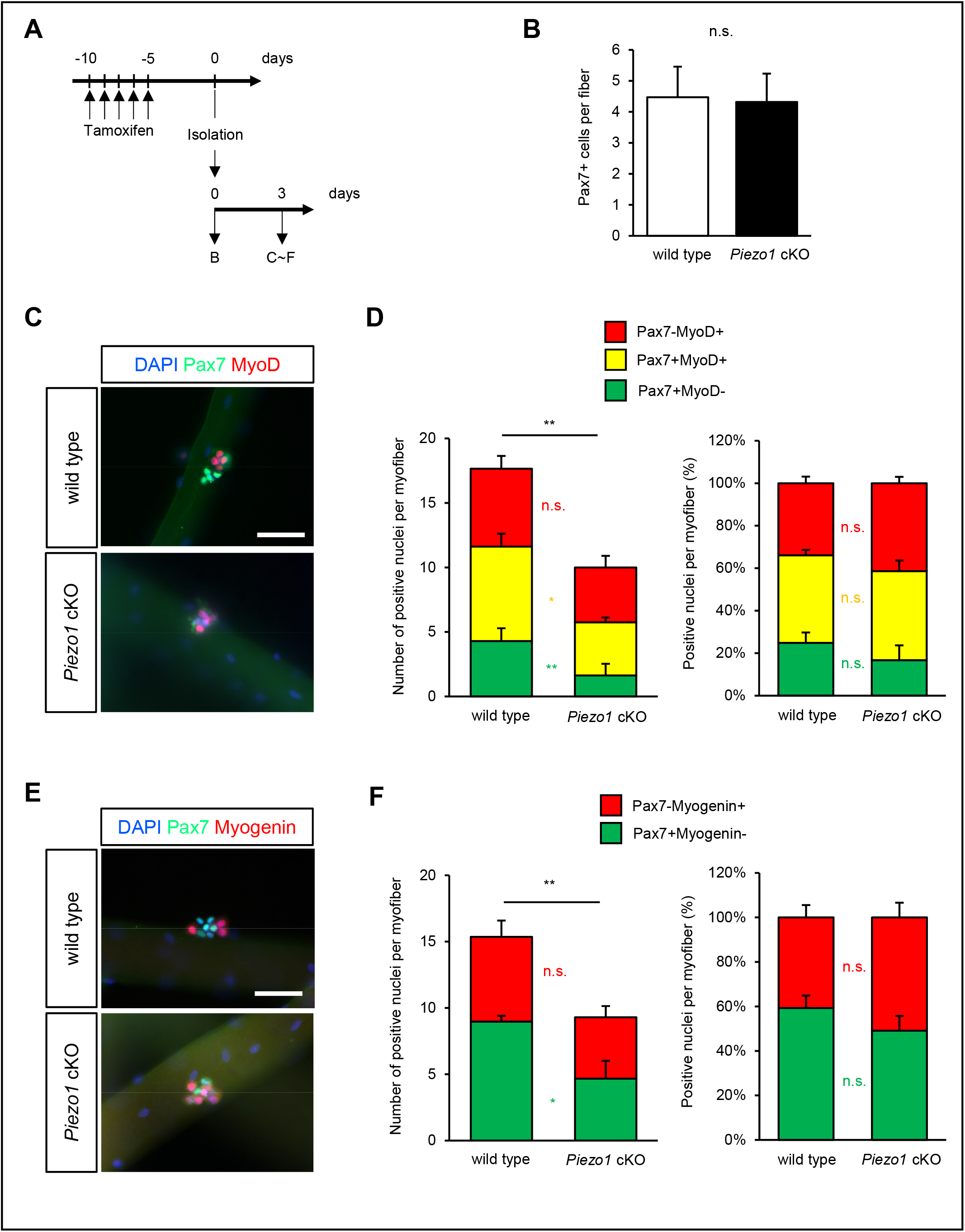
*Piezo1*-deficiency reduces the proliferative ability of MuSCs on isolated myofibers. **A** Time course for induction of *Piezo1*-deficiency and harvesting myofibers. **B** Detection of Pax7-positive MuSCs on freshly isolated myofibers (open column: wild type; closed column: *Piezo1* cKO MuSCs) (N = 5). **C** Immunofluorescent analysis of MuSCs on floating myofibers with anti-Pax7 and MyoD antibodies. Upper panel: wild type, lower panel: *Piezo1* cKO myofibers. Bar: 200 μm. **D** The number of Pax7-positive MuSCs per each myofiber, as shown in Fig 5C (N = 4). *: *P* < 0.05, **: *P* < 0.01. **E** Immunofluorescent analysis of MuSCs on floating myofibers with anti-Pax7 and myogenin antibodies. Upper panel: wild type, lower panel: *Piezo1* cKO myofibers. Bar: 200 μm. **F** Number of Pax7-positive MuSCs per myofiber, as shown in Fig 5E (N = 4).

### *Piezo1* deficiency affects the phosphorylation of myosin light chain in MuSCs

The changes in intracellular signaling for cytoskeletal reorganization are critical for the function and differentiation of MuSCs, including cell shape, cell migration, and maintenance of stemness [11]. Previous literatures have shown that a variety of intracellular signaling cascades are thought to be activated in a PIEZO1-dependent manner [25]. For example, we previously demonstrated that PIEZO1 activation is critical for RhoA/ROCK-mediated actomyosin formation in myoblasts, resulting in regulated myotube formation [26]. Thus, we hypothesized that PIEZO1 could act as an upstream activator for cytoskeleton rearrangement in MuSCs by phosphorylating myosin light chain (MLC) through the RhoA/ROCK pathway. To address this hypothesis, we performed immunofluorescent analysis to detect phosphorylated myosin light chain (pMLC) that is essential for the promotion of actomyosin formation [33]. The expression of pMLC was significantly reduced in *Piezo1* cKO compared to that in wild type mice (Figs 6A-6C). The same trend was observed in MuSCs seven days after cardiotoxin injection, where impaired regeneration capacity was observed in *Piezo1* cKO mice (EV5A and 5B). Importantly, treatment with CN03 (a RhoA activator) after MuSC activation on myofiber remarkably prevented the reduction in the MuSC number observed in *Piezo1* cKO mice (Fig 6D). Moreover, RhoA inhibition after MuSC activation on myofiber remarkably reduced the number of Pax7-positive cells (EV5C-5E). These results indicate that PIEZO1 plays a role in RhoA/ROCK-mediated phosphorylation of pMLC, which is critical for controlling the mechanical properties of MuSCs.

**Figure 6.**
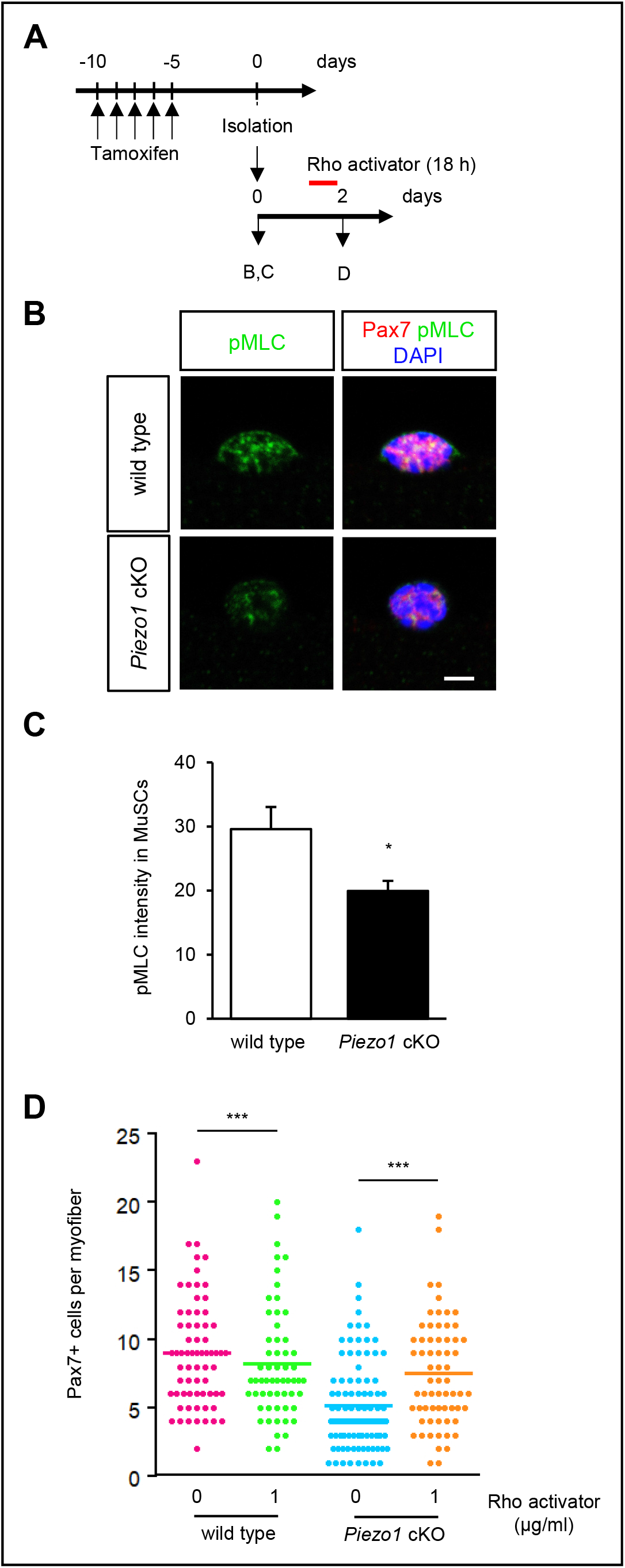
Reduction in phosphorylated MLC in *Piezo1*-deficient MuSCs. **A** Time course for the induction of *Piezo1*-deficiency, administration of RhoA activator, and harvesting myofibers. **B** Detection of phosphorylated MLC (pMLC) in MuSCs on myofibers from intact muscle. Pax7 and pMLC were detected with anti-Pax7 and pMLC antibodies, respectively. **C** Fluorescent intensity detected with anti-pMLC antibody was quantified, as shown in **b** (N = 5). Bar represents means ± S.E.M. **D** Effect of a RhoA activator CN03 on the proliferation of wild type or *Piezo1* cKO MuSCs (N = 7).

## Discussion

Since the population of MuSCs was identified in 1961 [1], great efforts have been made to understand the molecular mechanisms for MuSC-dependent muscle regeneration. Although the mechanical stimulation of MuSCs along with a series of secretory molecules have been thought to play fundamental roles in MuSC functions, the molecular entity that senses the changes in the mechanical properties of surrounding niche remains to be identified. In this study, we reported that PIEZO1 is involved in MuSC proliferation by mediating spontaneous Ca^2+^ influx across its plasma membrane. The deletion of *Piezo1* in MuSCs reduces its capacity to proliferate, leading to abnormalities in muscle regeneration. Thus, PIEZO1 plays a role in regenerating myofibers, probably by sensing the changes in the niche surrounding MuSCs.

Spontaneous Ca^2+^ influx is thought to be involved in a variety of physiological signals, especially in non-excitable cells [34]. Although mechanosensitive ionic currents were recorded first in chick myoblasts [35], the function of Ca^2+^ influx mediated by mechanosensation during myogenesis is poorly understood. Our results show that RhoA-mediated actomyosin formation occurs in a PIEZO1-dependent manner, suggesting that PIEZO1 acts as an upstream molecule for RhoA to control the mechanical properties of MuSCs. Previous studies have identified several candidate molecules that couple Ca^2+^ influx across the plasma membrane to RhoA-dependent actomyosin reorganization. For example, a Ca^2+^-dependent tyrosine kinase PYK2 phosphorylates PDZ-RhoGEF, one of Rho-specific guanine exchange factor (GEF), thereby activating RhoA in aortic vascular smooth muscle cells [36]. A Src family tyrosine kinase Fyn is another candidate molecule, as PIEZO2 is involved in activation of RhoA in breast cancer cells via the functional interaction between Fyn and one of GEFs LARG [37]. Moreover, Ca^2+^/calmodulin-dependent kinase (CaMKII) can also mediate Ca^2+^-dependent activation of RhoA in dendritic spines [38]. Further studies will be needed to elucidate the molecular mechanisms that underlie PIEZO1-mediated RhoA activation in MuSCs.

The changes in mechanical properties, such as actomyosin formation, are thought to be involved in MuSC functions. Recently, Eliazer et al. [11] demonstrated that myofiber-derived Wnt4 is essential for maintenance of stemness in MuSCs through RhoA-dependent repression of a transcription co-activator YAP1. We showed that PIEZO1 acts as the most upstream molecule; it may transduce mechanical signals into RhoA-mediated actomyosin formation to control the mechanical properties of activated MuSCs. Perhaps PIEZO1 and Wnt4-signaling via frizzled receptors might cooperatively enhance RhoA-mediated actomyosin formation, enabling MuSCs to maintain their functions.

The microenvironment is known to have a major impact on the growth and function of MuSCs [39]. Under physiological and pathological conditions, the mechanical properties of MuSC niche are thought to be changed; the stiffness of myofibers in regenerating muscle and in aged mice is greater than in normal and young mice, probably due to the changes in the composition and/or the amount of the extracellular matrix [40–42]. The importance of the MuSC niche is further confirmed by the fact that MuSCs cultured in substrates with elasticity mimicking the native myofibers retain the ability to self-renew or maintain quiescence compared to those cultured in conventional plastic dishes [43–45]. Despite these findings, how MuSCs itself sense the rigidity of the niche has not yet been elucidated. Given that PIEZO1 is essential for mechanosensation in a variety of cell types and tissues, PIEZO1 may also be the mechanosensor that plays a role in MuSC function.

Along with ion channels, a variety of membrane proteins have been identified as mechanosensors in physiological systems [46–48]. It is tempting to speculate that a series of mechanosensitive proteins have distinct roles in promoting the functions of MuSCs, such as its proliferation, differentiation, and migration. Further studies will unveil the role of mechanosensors in the progression and completion of myogenesis.

## Materials and Methods

### Mice

Animal care, ethical usage, and protocols were approved by the Animal Care Use and Review Committee of the Graduate School of Engineering, Kyoto University. Sperm samples for transgenic mouse strains, *Piezo1^tm1a(KOMP)Wtsi^* and *Piezo1^tm1c(KOMP)Wtsi^* [49], were purchased from the UC Davis KOMP repository. Cryo-recovery of the sperm was carried out by RIKEN BioResource Research Center (Japan). *Piezo1^tm1c(KOMP)Wtsi^* mice were further mated with *Pax7-*CreERT2/+ transgenic mice (The Jax laboratory, Strain ID: 012476; [29]) to generate MuSC-specific *Piezo1-*deficient mice. A *Piezo1-tdTomato* mouse line [27] was kindly provided from Dr. Keiko Nonomura (NIBB, Japan).

Tamoxifen (Sigma; dissolved in corn oil at a concentration of 20 mg/mL) was utilized to induce the expression of the Cre-recombinase. The mice were injected intraperitoneally with TMX at 40 μg/g of body weight for five consecutive days (in Figs 2, 3) and injected with 2 mg TMX daily for five days for ex vivo studies.

### MuSCs isolation using FACS

MuSCs from uninjured limb muscles were isolated as previously described [50]. Briefly, skeletal muscle samples taken from the forelimbs of mice were subjected to collagenase treatment using 0.2% collagenase type I (Sigma). Mononuclear cells were incubated with APC (allophycocyanin) or PE (phycoerythrin)-conjugated anti-mouse Ly-6A/E (Sca-1) antibody (#122508, BioLegend), APC or PE-conjugated anti-mouse CD45 antibody (#103106, BioLegend), APC or PE-conjugated anti-mouse CD31 antibody (#102508, BioLegend), and APC or PE-conjugated anti-mouse CD106 antibody (#105718, BioLegend) at 4°C for at least 30 min. Those cells were resuspended in PBS containing 2% FBS, and then subjected to cell sorting to collect CD106-positive cells using MA900 (Sony).

### RT-PCR

For RT-PCR analysis, MuSCs (QSCs and ASCs) and gastrocnemius muscle were isolated from Pax7-YFP mice [50] and C57BL/6J mice, respectively. Total RNA was isolated using ISOGEN II (Nippon Gene) or Qiagen RNeasy Micro Kit. cDNA was generated with PrimeScript™ II 1st strand cDNA Synthesis Kit (Takara). Semi-quantitative PCR was performed with EmeraldAmp MAX PCR Master Mix (Takara). Primers for amlplication of *Piezo1* and *Gapdh* genes were listed in our previous study [26].

### Ca^2+^ imaging in MuSCs

Fura-2 imaging was performed as described previously [26]. For indicator loading, MuSCs were plated on glass-bottomed dishes (Matsunami) coated with Matrigel, then incubated with 5 μM Fura-2 AM (Dojindo) at 37°C for 60 min. Time-lapse images were obtained at every 2s. Ratiometric images (F340/F380) were analyzed with a physiology software (Zeiss). Yoda1-induced Ca^2+^ influx was quantified as the difference in the Fura-2 ratio between its maximum value and at 1 min from imaging initiation. These experiments were carried out using a heat chamber (Zeiss) to maintain 37°C throughout the imaging.

### Single myofiber isolation

Myofibers were isolated from EDL muscles, as described previously [51]. Isolated EDL muscle samples were incubated with 0.2% collagenase I (sigma) in DMEM at 37°C for 2 hours. Myofibers were released by gently flushing the muscle samples in the plating medium using a fire-polished glass pipette.

### Cell culture

MuSCs were cultured in a growth medium (DMEM supplemented with 30% fetal bovine serum, 1% chick embryo extract (US Biological), 10 ng/mL basic fibroblast growth factor (ORIENTAL YEAST Co., Ltd.), and 1% penicillin-streptomycin) on culture dishes coated with Matrigel (Corning). 4-OH TMX (Sigma) (1 μM) was added to both wild type and *Piezo1* cKO growth medium for freshly isolated MuSC culture. Twenty μM of BAPTA-AM (Dojindo) was added to Growth medium during culture. Ten μM of EdU (Life Technologies) was added to the growth medium during culture.

For MuSC growth on floating myofibers, isolated myofibers were cultured in a plating medium (DMEM supplemented with 10% horse serum, 0.5% chick embryo extract, and 1% penicillin-streptomycin) at 37°C with 5% CO_2_. Rho activator II (CN03, Cytoskeleton, Inc.) or Rho inhibitor I (CT04, Cytoskeleton, Inc.) were added to the plating medium 30 h after plating.

### Immunofluorescent analysis

The cells were placed onto Matrigel-coated glass bottom dishes and fixed with 4% paraformaldehyde (PFA)/PBS for 10 min. The myofibers were fixed with 2% PFA/PBS for 5 to 10 min. After permeabilization in 0.1% Triton X-100/PBS for 15 min, those samples were blocked in 1% BSA/PBS for 30 min and probed with the following antibodies at room temperature for 2 hours or at 4℃ overnight: anti-PAX7 antibody (Developmental Studies Hybridoma Bank, mouse; 1:500), anti-RFP antibody (600-401-379, Rockland, rabbit; 1:500), anti-MyoD antibody (sc-304, Santa Cruz, rabbit; 1:1000), anti-Myogenin (sc-576, Santa Cruz, rabbit; 1:500), and anti-P-MLC antibody (3671, CST, rabbit; 1:200). Nuclei were detected using DAPI (Dojindo, 1:1000). Immunofluorescent signals were visualized with Alexa488- or Alexa555-conjugated secondary antibodies, using a confocal microscope (LSM 800, Zeiss) with a 63x objective lens. Fluorescence intensity was quantified using ImageJ software for statistical analyses.

For EdU detection, the click chemical reaction was performed after the primary and secondary staining using a Click-iT EdU Imaging Kit (Life Technologies) according to the manufacturer’s instructions.

### Histological analysis

Cardiotoxin experiments were carried out according to previous literature [26]. Fifty μL of 10 μM cardiotoxin (Latoxan) was injected into the TA muscle of 8-to 15-week-old mice. The muscle was harvested at time points as indicated in each figure, and snap-frozen in isopentane cooled with liquid nitrogen. The cross cryosections (thickness, 7 μm) of the muscle samples were used for Hematoxylin & Eosin staining, as previously described [52]. CSA in each muscle sample was determined on cryosections stained with anti-Laminin I antibody (L9393, Sigma Aldrich 1:500). Fibrotic area was detected using anti-Collagen I antibody (1310-01, Southern Biotech, 1:500). CSA and fluorescence intensity were quantified using ImageJ software for statistical analyses.

### In vivo EdU-uptake assay

EdU was dissolved in PBS at 0.5 mg/mL and injected intraperitoneally at 0.1 mg per 20 g body weight at time points indicated in each figure.

## Acknowledgments

We thank Drs. Hiroshi Takeshima and Atsuhiko Ichimura (Kyoto University) for helpful discussion. We also thank Dr. Itaru Hamachi (Kyoto University) and all the lab members for technical support. This study was supported by the Japan Agency for Medical Research and Development (AMED, JP19gm5810016); the Grant-in-Aid for Scientific Research KAKENHI (19H03179); Intramural Research Grant (2-5) for Neurological and Psychiatric Disorder of NCNP; grants from the Takeda science foundation, Ohsumi Frontier Science Foundation, Nakatomi Foundation, and Asahi Glass Foundation (to Y.H.); and Grant-in-Aid for JSPS Research Fellow (20J22984 to K.H.)

## Author contributions

K.H., M.T. S.T., K.Na., M.U., and Y.H. designed the project. K.H., and Y.H. interpreted data and wrote the paper. K.H. performed the experiments. M.T. established FACS system for MuSC isolation. Y.K. and Y.O. supervised assays to evaluate MuSC functions. K.No., and Y.M. supervised PIEZO1 analysis.

## Expanded View Figure legends

**Figure EV1.**
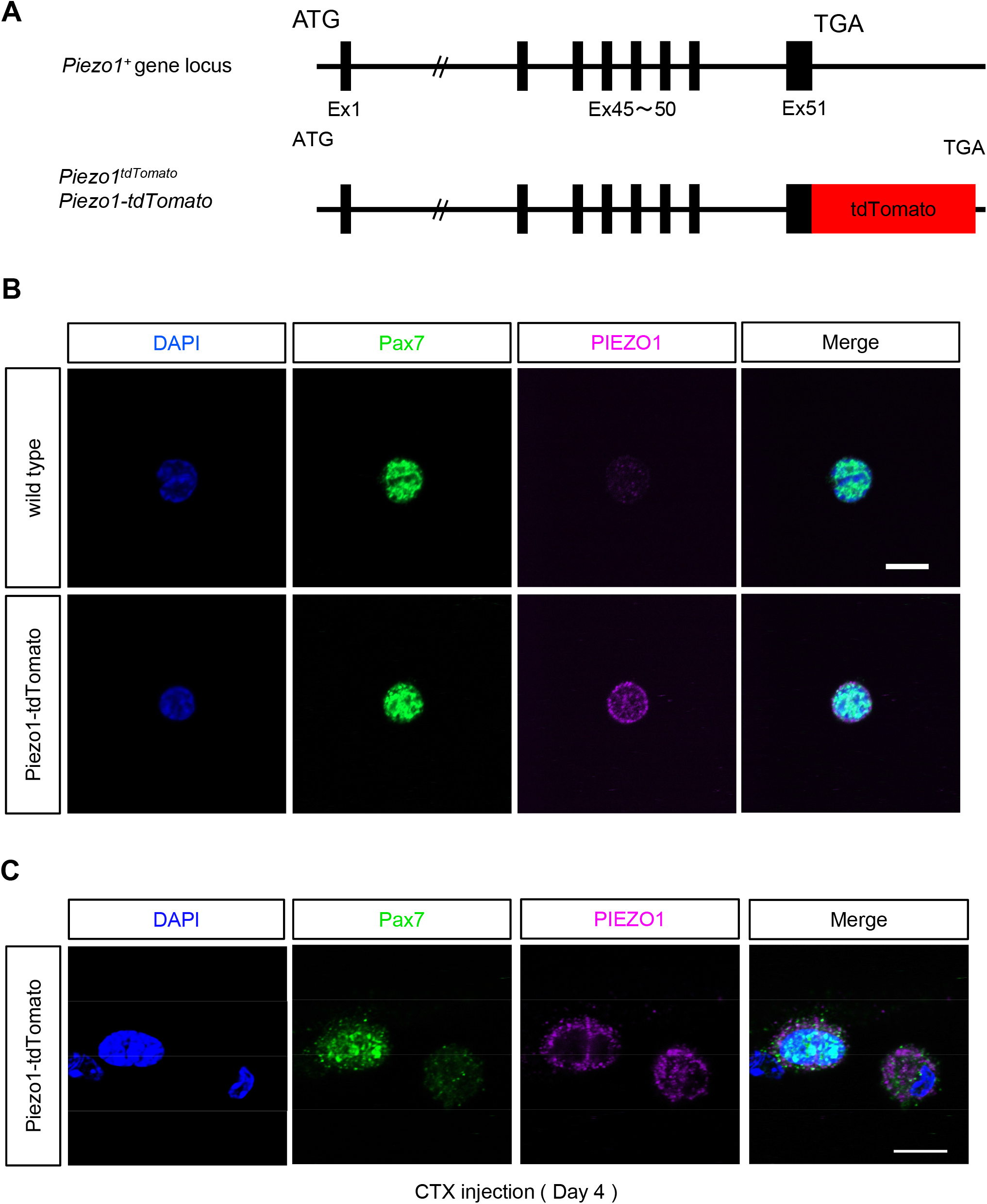
Expression of PIEZO1-tdTomato in MuSCs. **A** Schematic figure representing the *Piezo1* locus in *Piezo1-tdTomato* mice. **B** Expression of PIEZO1-tdTomato in isolated MuSCs. MuSCs were isolated from *Piezo1-tdTomato* and wild type muscle tissues, and then stained for tdTomato and PAX7. Upper panels: MuSCs from wild type; lower panels: PIEZO1-tdTomato. **C** Detection of PIEZO1-tdTomato in regenerating myofibers. Myofibers were isolated from cardiotoxin-injected EDL muscles four days post-injection.

**Figure EV2.**
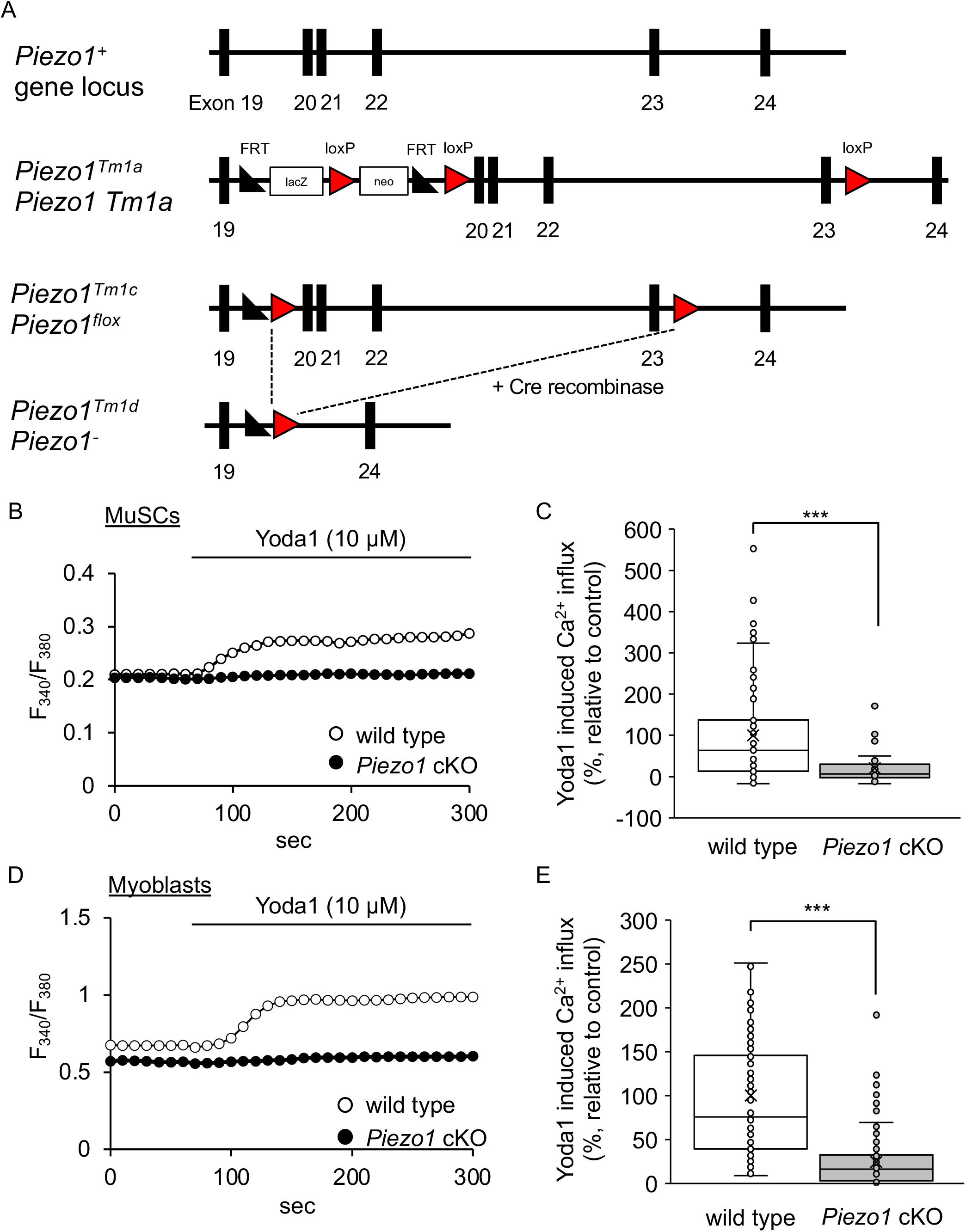
Generation of MuSC-specific *Piezo1*-deficient mice. **A** Schematic figure representing the *Piezo1* locus in *Piezo1* cKO mice. **B, C** Ca^2+^ measurements in MuSCs and myoblasts isolated from *Piezo1* cKO mice. Endogenous PIEZO1 in MuSCs (**B**) or myoblasts (**C**) was activated by the administration of a PIEZO1 agonist, Yoda1.

**Figure EV3.**
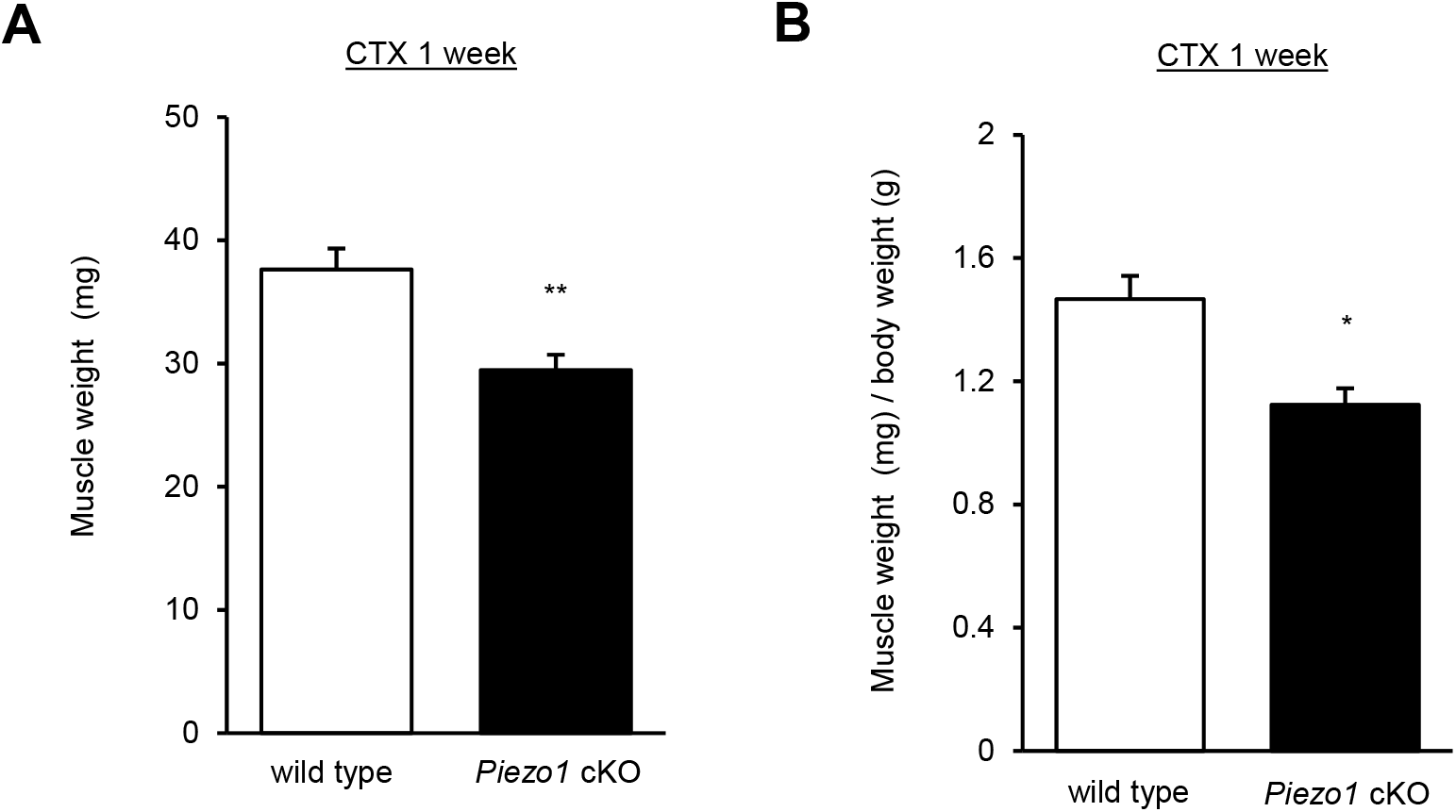
Measurements of muscle weight isolated from wild type and *Piezo1* cKO mice one week post-cardiotoxin injection. **A** Muscle (TA) weight of wild type or *Piezo1* cKO mice. N = 3–6 **B** Muscle weight / body weight of wild type or *Piezo1* cKO mice. N = 3–6

**Figure EV4.**
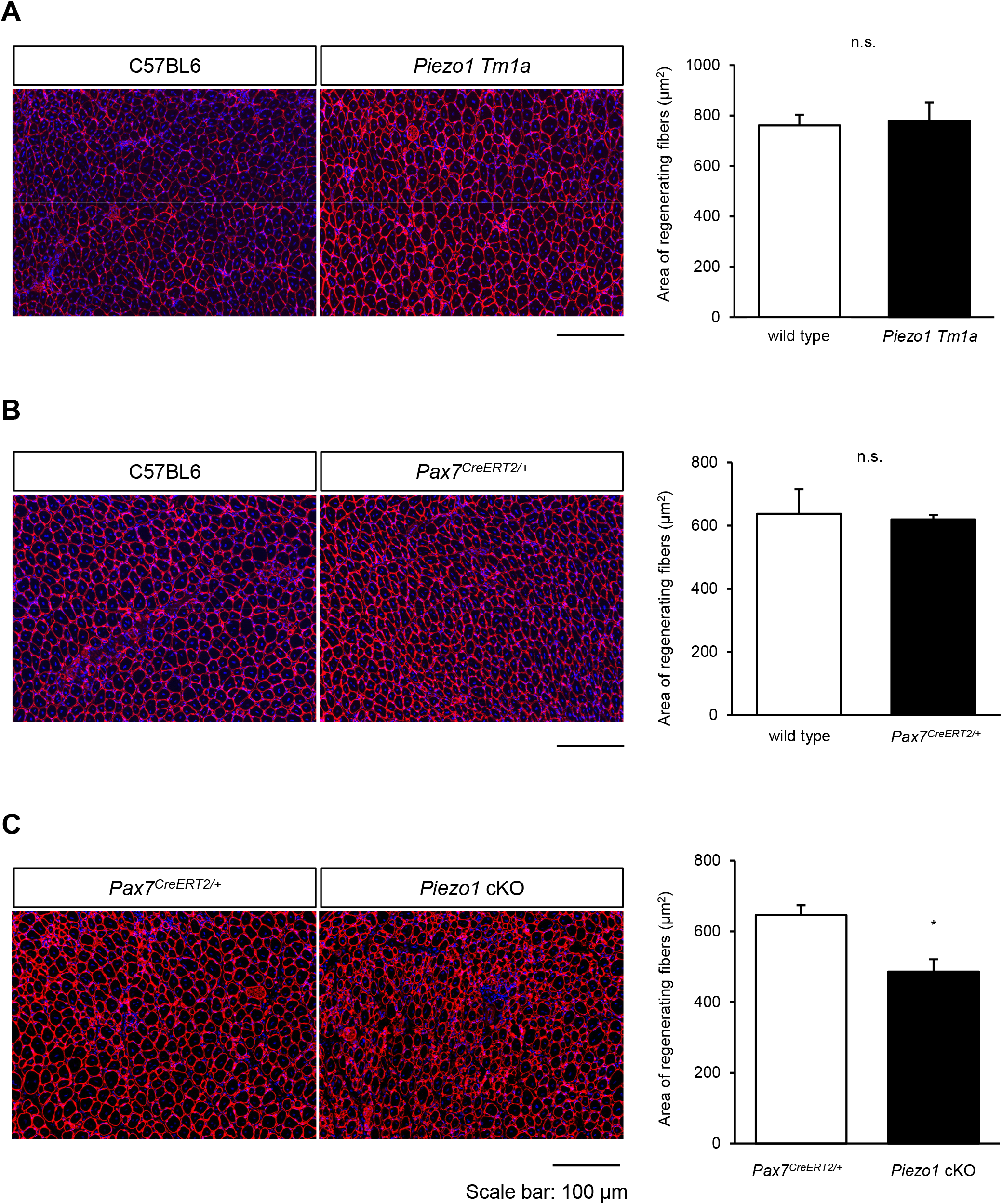
Evaluation of regenerating myofibers in *Piezo1*-heterozygous mice. **A**-**C** Evaluation of Cross Section Area (CSA μm^2^) in *Piezo1 Tm1a* (**A**), *Piezo1^+/+^*; *Pax7^CreERT2/+^* (**B**), and *Piezo1* cKO (**C**) muscle one week post-cardiotoxin injection. Cross sections from those mice were stained with anti-laminin antibody and DAPI, and then evaluated for CSA. Bar: 100 μm.

**Figure EV5.**
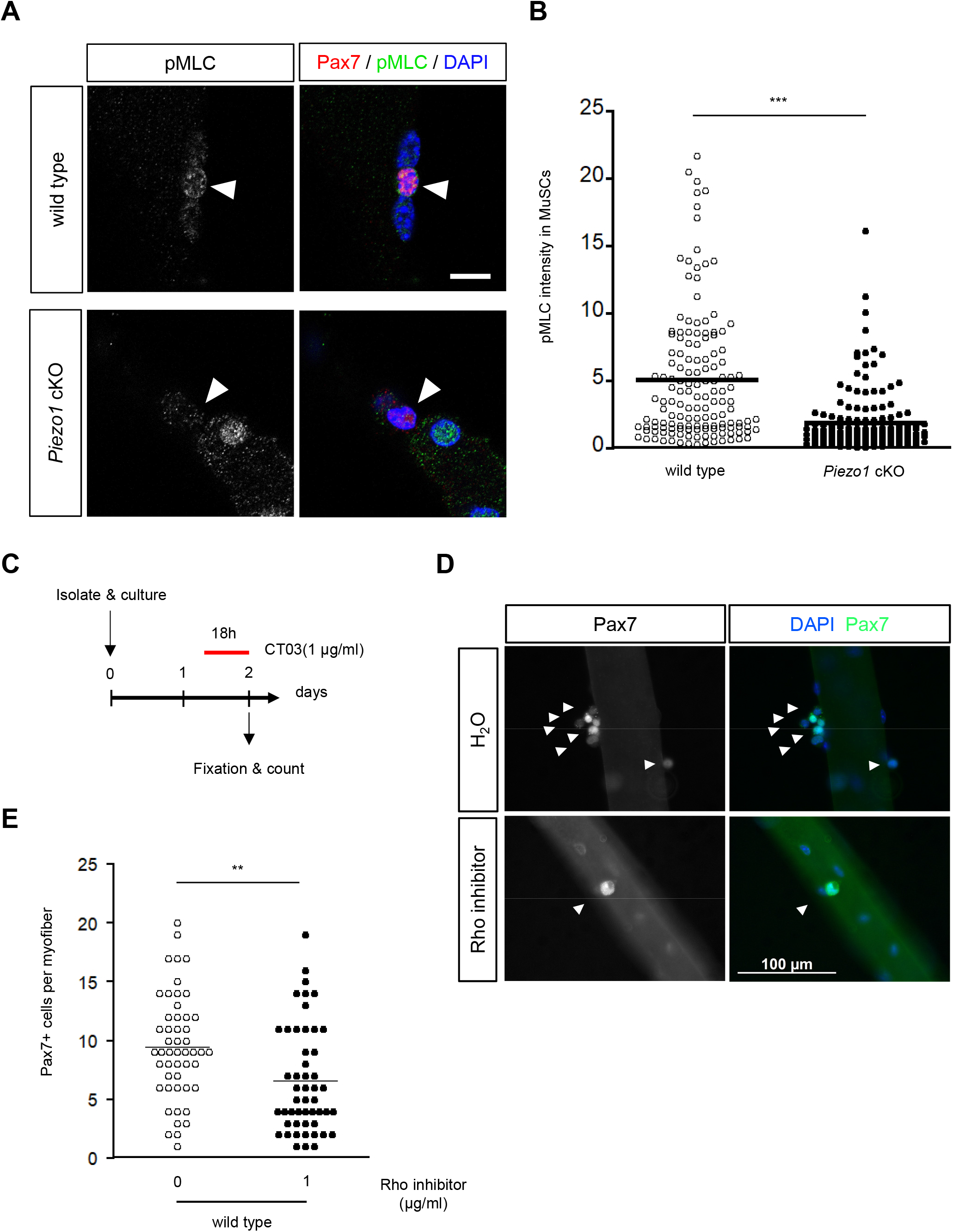
Detection of phosphorylated form of MLC in MuSCs from regenerating muscle tissues. **A, B** Myofibers were isolated from EDL one week post-cardiotoxin injection and immunostained with anti-Pax7 and anti-pMLC antibody. **C** Time course of RhoA inhibition during MuSC proliferation on myofiber. **D** Immunofluorescent analysis of MuSCs on floating myofibers with anti-Pax7 after RhoA inhibition. **E** Effect of a RhoA inhibitor on the proliferation of wild type. (N = 3)

## References

1. Mauro, A., Satellite cell of skeletal muscle fibers. J Biophys Biochem Cytol, 1961. 9: p. 493–5.

2. Relaix, F. and P.S. Zammit, Satellite cells are essential for skeletal muscle regeneration: the cell on the edge returns centre stage. Development, 2012. 139(16): p. 2845–56.

3. Dumont, N.A., et al., Dystrophin expression in muscle stem cells regulates their polarity and asymmetric division. Nat Med, 2015. 21(12): p. 1455–63.

4. Tierney, M.T. and A. Sacco, Satellite Cell Heterogeneity in Skeletal Muscle Homeostasis. Trends Cell Biol, 2016. 26(6): p. 434–444.

5. Feige, P., et al., Orienting Muscle Stem Cells for Regeneration in Homeostasis, Aging, and Disease. Cell Stem Cell, 2018. 23(5): p. 653–664.

6. Seale, P., et al., Pax7 is required for the specification of myogenic satellite cells. Cell, 2000. 102(6): p. 777–786.

7. Troy, A., et al., Coordination of satellite cell activation and self-renewal by Par-complex-dependent asymmetric activation of p38alpha/beta MAPK. Cell Stem Cell, 2012. 11(4): p. 541–53.

8. Bjornson, C.R., et al., Notch signaling is necessary to maintain quiescence in adult muscle stem cells. Stem Cells, 2012. 30(2): p. 232–42.

9. Fujimaki, S., et al., Notch1 and Notch2 Coordinately Regulate Stem Cell Function in the Quiescent and Activated States of Muscle Satellite Cells. Stem Cells, 2018. 36(2): p. 278–285.

10. Mourikis, P., et al., A critical requirement for notch signaling in maintenance of the quiescent skeletal muscle stem cell state. Stem Cells, 2012. 30(2): p. 243–52.

11. Eliazer, S., et al., Wnt4 from the Niche Controls the Mechano-Properties and Quiescent State of Muscle Stem Cells. Cell Stem Cell, 2019. 25(5): p. 654–665 e4.

12. Rodgers, J.T., et al., HGFA Is an Injury-Regulated Systemic Factor that Induces the Transition of Stem Cells into GAlert. Cell Rep, 2017. 19(3): p. 479–486.

13. Clapham, D.E., Calcium signaling. Cell, 2007. 131(6): p. 1047–58.

14. Berridge, M.J., M.D. Bootman, and H.L. Roderick, Calcium signalling: dynamics, homeostasis and remodelling. Nat Rev Mol Cell Biol, 2003. 4(7): p. 517–29.

15. Calderon, J.C., P. Bolanos, and C. Caputo, The excitation-contraction coupling mechanism in skeletal muscle. Biophys Rev, 2014. 6(1): p. 133–160.

16. Campbell, K.P., A.T. Leung, and A.H. Sharp, The biochemistry and molecular biology of the dihydropyridine-sensitive calcium channel. Trends in Neurosciences, 1988. 11(10): p. 425–430.

17. Michelucci, A., et al., Role of STIM1/ORAI1-mediated store-operated Ca(2+) entry in skeletal muscle physiology and disease. Cell Calcium, 2018. 76: p. 101–115.

18. Endo, Y., et al., Dominant mutations in ORAI1 cause tubular aggregate myopathy with hypocalcemia via constitutive activation of store-operated Ca(2)(+) channels. Hum Mol Genet, 2015. 24(3): p. 637–48.

19. Coste, B., et al., Piezo1 and Piezo2 are essential components of distinct mechanically activated cation channels. Science, 2010. 330(6000): p. 55–60.

20. Ma, S., et al., Common PIEZO1 Allele in African Populations Causes RBC Dehydration and Attenuates Plasmodium Infection. Cell, 2018. 173(2): p. 443–455 e12.

21. Pathak, M.M., et al., Stretch-activated ion channel Piezo1 directs lineage choice in human neural stem cells. Proc Natl Acad Sci U S A, 2014. 111(45): p. 16148–53.

22. Lee, W., et al., Synergy between Piezo1 and Piezo2 channels confers high-strain mechanosensitivity to articular cartilage. Proc Natl Acad Sci U S A, 2014. 111(47): p. E5114–22.

23. Li, J., et al., Piezo1 integration of vascular architecture with physiological force. Nature, 2014. 515(7526): p. 279–282.

24. Nonomura, K., et al., Mechanically activated ion channel PIEZO1 is required for lymphatic valve formation. Proc Natl Acad Sci U S A, 2018. 115(50): p. 12817–12822.

25. Murthy, S.E., A.E. Dubin, and A. Patapoutian, Piezos thrive under pressure: mechanically activated ion channels in health and disease. Nat Rev Mol Cell Biol, 2017. 18(12): p. 771–783.

26. Tsuchiya, M., et al., Cell surface flip-flop of phosphatidylserine is critical for PIEZO1-mediated myotube formation. Nat Commun, 2018. 9(1): p. 2049.

27. Ranade, S.S., et al., Piezo1, a mechanically activated ion channel, is required for vascular development in mice. Proc Natl Acad Sci U S A, 2014. 111(28): p. 10347–52.

28. Yin, H., F. Price, and M.A. Rudnicki, Satellite cells and the muscle stem cell niche. Physiol Rev, 2013. 93(1): p. 23–67.

29. Lepper, C. and C.M. Fan, Inducible lineage tracing of Pax7-descendant cells reveals embryonic origin of adult satellite cells. Genesis, 2010. 48(7): p. 424–36.

30. Syeda, R., et al., Chemical activation of the mechanotransduction channel Piezo1. Elife, 2015. 4.

31. Zammit, P.S., et al., Muscle satellite cells adopt divergent fates: a mechanism for self-renewal? J Cell Biol, 2004. 166(3): p. 347–57.

32. Ono, Y., et al., BMP signalling permits population expansion by preventing premature myogenic differentiation in muscle satellite cells. Cell Death Differ, 2011. 18(2): p. 222–34.

33. Vicente-Manzanares, M., D.J. Webb, and A.R. Horwitz, Cell migration at a glance. J Cell Sci, 2005. 118(Pt 21): p. 4917–9.

34. Qian, N., et al., TRPM7 channels mediate spontaneous Ca(2+) fluctuations in growth plate chondrocytes that promote bone development. Sci Signal, 2019. 12(576).

35. Guharay, F. and F. Sachs, Stretch-activated single ion channel currents in tissue-cultured embryonic chick skeletal muscle. J Physiol, 1984. 352: p. 685–701.

36. Ying, Z., et al., PYK2/PDZ-RhoGEF links Ca2+ signaling to RhoA. Arterioscler Thromb Vasc Biol, 2009. 29(10): p. 1657–63.

37. Pardo-Pastor, C., et al., Piezo2 channel regulates RhoA and actin cytoskeleton to promote cell mechanobiological responses. Proc Natl Acad Sci U S A, 2018. 115(8): p. 1925–1930.

38. Murakoshi, H., H. Wang, and R. Yasuda, Local, persistent activation of Rho GTPases during plasticity of single dendritic spines. Nature, 2011. 472(7341): p. 100–4.

39. Fuchs, E. and H.M. Blau, Tissue Stem Cells: Architects of Their Niches. Cell Stem Cell, 2020. 27(4): p. 532–556.

40. Urciuolo, A., et al., Collagen VI regulates satellite cell self-renewal and muscle regeneration. Nat Commun, 2013. 4: p. 1964.

41. Trensz, F., et al., Increased microenvironment stiffness in damaged myofibers promotes myogenic progenitor cell proliferation. Skelet Muscle, 2015. 5: p. 5.

42. Lacraz, G., et al., Increased Stiffness in Aged Skeletal Muscle Impairs Muscle Progenitor Cell Proliferative Activity. PLoS One, 2015. 10(8): p. e0136217.

43. Cosgrove, B.D., et al., Rejuvenation of the muscle stem cell population restores strength to injured aged muscles. Nat Med, 2014. 20(3): p. 255–64.

44. Gilbert, P.M., et al., Substrate elasticity regulates skeletal muscle stem cell self-renewal in culture. Science, 2010. 329(5995): p. 1078–81.

45. Quarta, M., et al., An artificial niche preserves the quiescence of muscle stem cells and enhances their therapeutic efficacy. Nat Biotechnol, 2016. 34(7): p. 752–9.

46. Xu, J., et al., GPR68 Senses Flow and Is Essential for Vascular Physiology. Cell, 2018. 173(3): p. 762–775 e16.

47. Kefauver, J.M., A.B. Ward, and A. Patapoutian, Discoveries in structure and physiology of mechanically activated ion channels. Nature, 2020. 587(7835): p. 567–576.

48. Vining, K.H. and D.J. Mooney, Mechanical forces direct stem cell behaviour in development and regeneration. Nat Rev Mol Cell Biol, 2017. 18(12): p. 728–742.

49. West, D.B., et al., A lacZ reporter gene expression atlas for 313 adult KOMP mutant mouse lines. Genome Res, 2015. 25(4): p. 598–607.

50. Kitajima, Y. and Y. Ono, Visualization of PAX7 protein dynamics in muscle satellite cells in a YFP knock-in-mouse line. Skelet Muscle, 2018. 8(1): p. 26.

51. Nagata, Y., et al., Sphingomyelin levels in the plasma membrane correlate with the activation state of muscle satellite cells. J Histochem Cytochem, 2006. 54(4): p. 375–84.

52. Hara, Y., et al., A dystroglycan mutation associated with limb-girdle muscular dystrophy. N Engl J Med, 2011. 364(10): p. 939–46.

